# Single source of pangolin CoVs with a near identical Spike RBD to SARS-CoV-2

**DOI:** 10.1101/2020.07.07.184374

**Authors:** Yujia Alina Chan, Shing Hei Zhan

**Author notes:** Authors contributed equally to this work.

## Abstract

Multiple publications have independently described pangolin CoV genomes from the same batch of smuggled pangolins confiscated in Guangdong province in March, 2019. We analyzed the three metagenomic datasets that sampled this batch of pangolins and found that the two complete pangolin CoV genomes, GD_1 by Xiao et al. *Nature* and MP789 by Liu et al. *PLoS Pathogens*, were both built primarily using the 2019 dataset first described by Liu et al. *Viruses*. Other publications, such as Zhang et al. *Current Biology* and Lam et al. *Nature*, have also relied on this same dataset by Liu et al. *Viruses* for their assembly of the Guangdong pangolin CoV sequences and comparisons to SARS-CoV-2. To our knowledge, all of the published pangolin CoV genome sequences that share a highly similar Spike receptor binding domain with SARS-CoV-2 originate from this singular batch of smuggled pangolins. This raises the question of whether pangolins are truly reservoirs or hosts of SARS-CoV-2-related coronaviruses in the wild, or whether the pangolins may have contracted the CoV from another host species during trafficking. Our observations highlight the importance of requiring authors to publish their complete genome assembly pipeline and all contributing raw sequence data, particularly those supporting epidemiological investigations, in order to empower peer review and independent analysis of the sequence data. This is necessary to ensure both the accuracy of the data and the conclusions presented by each publication.

---

Several studies describing pangolin CoV genomes have posited that pangolins are potentially important hosts in the emergence of novel coronaviruses. There has been particular emphasis on the discovery of a Guangdong pangolin CoV that shares all five critical residues and ~97% amino acid identity in the Spike receptor binding domain (RBD) with SARS-CoV-2.^1–3^ This is due to the fact that the Spike RBD plays a major role in determining the host specificity of each coronavirus. The next closest match to SARS-CoV-2 in terms of the Spike RBD is the bat CoV RaTG13, sharing 90.13% Spike RBD amino acid identity. RaTG13 shares the highest genome identity to SARS-CoV-2 (96%), whereas the Guangdong pangolin CoV shares ~90% genome identity with SARS-CoV-2. Nonetheless, based on Spike RBD similarity, many have hypothesized that pangolins could have been an intermediate host of SARS-CoV-2 that later transmitted into humans.

The Guangdong pangolin CoV, obtained from a batch of 21 smuggled pangolins in Guangdong province in March, 2019, was originally described in October, 2019 by Liu et al. *Viruses*.^4^ Since then, Liu et al. *PLoS Pathogens*,^1^ Xiao et al. *Nature*,^2^ Zhang et al. *Current Biology*,^3^ and Lam et al. *Nature*^5^ have each analyzed CoV sequences from this same batch of pangolins. In addition, Lam et al. describe CoVs from another batch of smuggled pangolins in Guangxi province. However, these Guangxi pangolin CoVs do not share such a high level of Spike RBD identity (only 86-87% amino acid identity) with SARS-CoV-2. The Guangdong pangolin CoV sequences have already been utilized across numerous other studies. Therefore, we analyzed the contributions of individual sequences from each study to identify the most representative Guangdong pangolin CoV genome for use by the research community.

During this process, we were surprised to find that Xiao et al. assembled their genome GD_1 (GISAID: EPI_ISL_410721) primarily using the Liu et al. *Viruses* (2019) metagenomic dataset (NCBI SRA BioProject: PRJNA573298) (**Fig 1, Supplementary Fig 1**). In their paper, Xiao et al. identified the PRJNA573298 dataset as a source of contigs from pangolin viral metagenomes that mapped to the SARS-CoV-2 genome. Yet, Xiao et al. cite Liu et al. *Viruses* in their article as follows: “*SARSr-CoV sequences were previously detected in dead Malayan pangolins (reference 15: Liu et al. Viruses). These sequences appear to be from Pangolin-CoV identified in the present study judged by their sequence similarity*.” Given that Xiao et al. relied predominantly on the dataset by Liu et al. *Viruses* to generate their genome sequence, it is curious that it was not clear to Xiao et al. that they were studying the same pangolin CoV.

**Figure 1.**
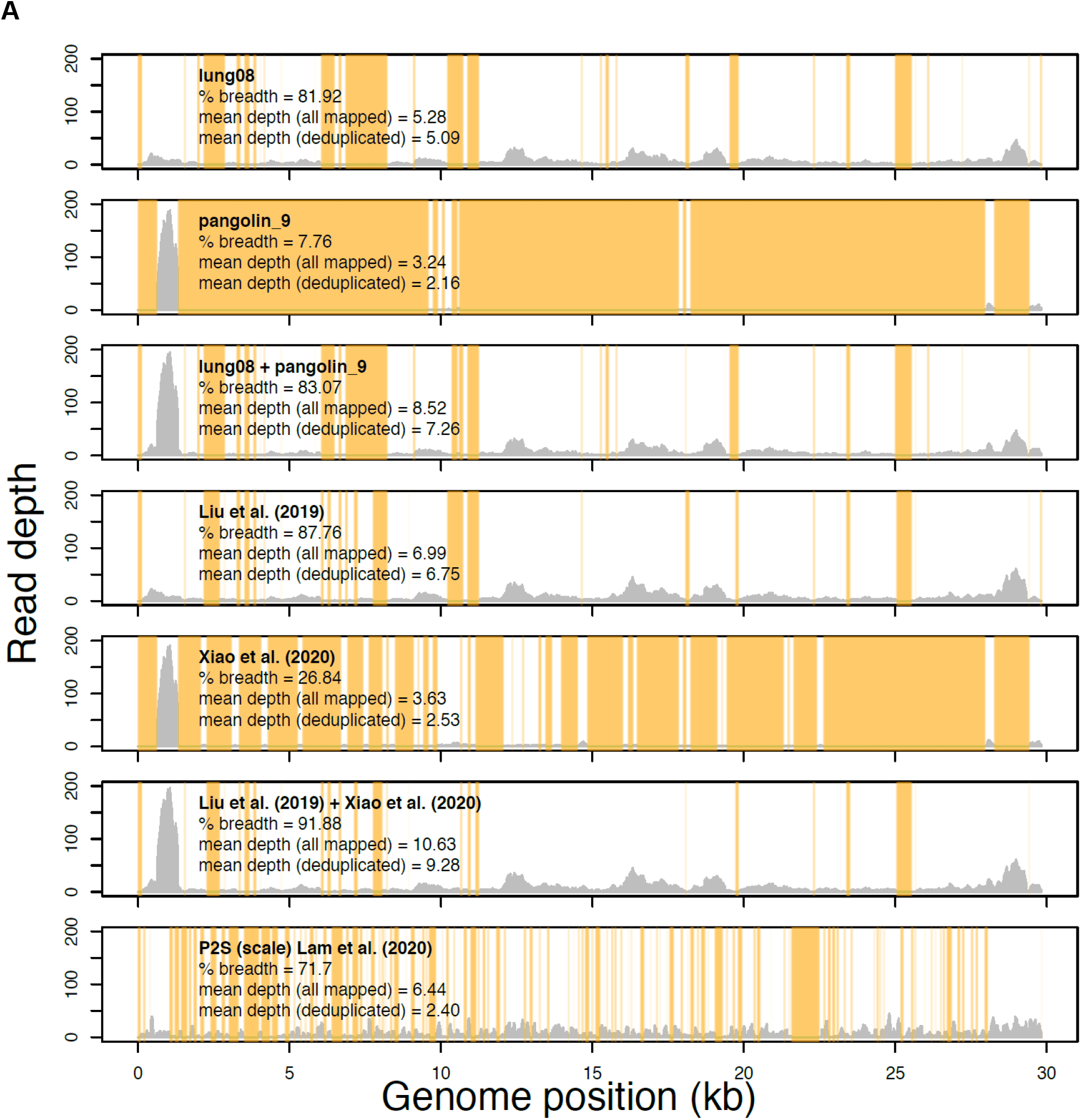
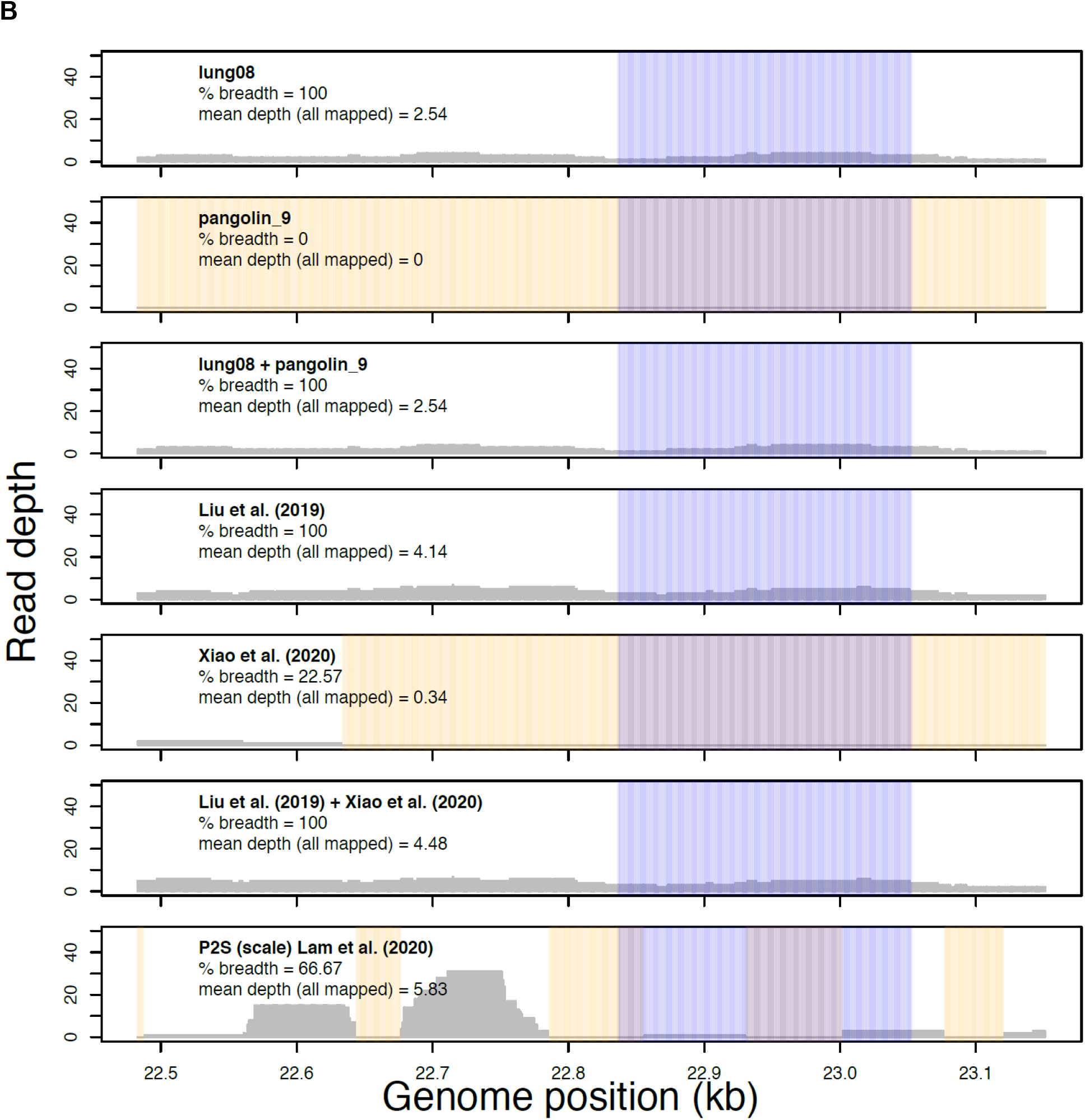
Read profiles of the metagenomic data sets from Liu et al. 2019 *Viruses*, Xiao et al. 2020 *Nature*, and Lam et al. 2020 *Nature* mapped to the Xiao et al. Guangdong pangolin CoV genome sequence GD_1 (EPI_ISL_410721). Samples lung08 (described in Liu et al. *Viruses* but re-introduced as M4 by Xiao et al. *Nature*) and pangolin_9 (sample M1, Xiao et al. *Nature*) each had the most sequence data of all samples analyzed in Liu et al. *Viruses* and Xiao et al. *Nature*, respectively. The “lung08 + pangolin_9” track shows their combined read coverage. The “Liu et al. (2019)” track indicates the read coverage pooled from all of the pangolin samples with mapped reads. The “Xiao et al. (2020)” track reveals the read coverage pooled from all samples unique to Xiao et al. *Nature* with mapped reads (**Supplementary Table 1**). The “Liu (2019) + Xiao (2020)” track combines the read coverage from both studies and appears highly similar to the metagenomic data reported in Xiao et al. *Nature* (**Supplementary Fig 1**). The “P2S Lam et al. (2020)” track shows the read coverage from the scale sample that was sequenced only by Lam et al. *Nature* (a single-end library). Read profiles are shown over the entire genome (**A**) or the Spike receptor binding domain with the receptor binding motif highlighted in blue (**B**). Regions with zero read coverage are highlighted in orange. Also shown are the estimates of the mean depth of read coverage with or without duplicate reads (“all mapped” or “deduplicated”).

Xiao et al. renamed pangolin samples first published by Liu et al. *Viruses* without citing their study as the original article that described these samples, and used the metagenomic data from these samples in their analysis. To compound this issue, Xiao et al. used an inconsistent definition of “total reads” in the description of the samples in their Extended Data Table 3, which made it challenging to match the samples with those from Liu et al. *Viruses*. For samples M1 and M6, Xiao et al. counted the number of sequenced *fragments* in the library, which corresponds to their stated library size (**Supplementary Table 1**). For the remaining samples, Xiao et al. counted the number of sequenced *reads* (two reads per fragment in paired-end Illumina libraries; **Supplementary Table 1** shows that the “total reads” for these samples are double the library size). We were only able to match the pangolin samples from Liu et al. *Viruses* with those used in Xiao et al.’s analysis after becoming aware of this inconsistency. Samples lung02, lung07, lung08, and lung11 published by Liu et al. *Viruses* were re-named as M3, M2, M4, and M8, respectively, in Xiao et al. *Nature* (**Supplementary Table 1**).

The sample lung08, first analyzed by Liu et al. *Viruses*, was a major source of metagenomic reads spanning the entire pangolin CoV genome (**Fig 1**, **Supplementary Fig 1**). The only sample from Xiao et al. that we could not match to those from the Liu et al. dataset, which contributed a considerable number of reads was pangolin_9 (sample M1). However, the majority of the pangolin_9/M1 reads map onto a single region in Orf1a (**Fig 1A**, **Supplementary Fig 1**). Indeed, when the reads from samples lung08 and pangolin_9 are combined, the resulting read coverage and alignment profile looks highly similar to the sample read profile in Xiao et al.’s Extended Data Fig 4 (**Supplementary Fig 1**). Xiao et al. did not specify which of their seven samples with CoV sequence reads was described by this figure.^2^ In addition, for the Spike RBD, the majority of the metagenomic sequence data are derived from lung08 (even so, the read depth was low; **Fig 1B**).

During the review of this manuscript, on June 22, 2020, Xiao et al. added a new sample, pangolin_10 to their *Nature* paper’s bioproject (NCBI SRA BioProject PRJNA607174).^6^ This new sample was not described in the main text, figures, tables, or extended materials of the *Nature* paper that was published on May 7, 2020. Specifically, Xiao et al. stated in their main text that “Illumina RNAseq was used to identify viruses in the lung from nine pangolins.” All nine pangolins are accounted for in their Extended Data Table 3, albeit four were renamed (and unattributed) samples from the 2019 Liu et al. *Viruses* publication. Despite the addition of pangolin_10, it remains unclear as to which sample produced the read profile shown in their Extended Data Fig 4. Based on the sample read profile, it is very likely that the first public description of pangolin_10 can instead be found in their recent Li et al. bioRxiv preprint, which was also released on June 22, 2020, but with no attribution of the sample to the *Nature* paper or the bioproject into which they deposited the pangolin_10 data.^7^

To add to the challenge of evaluating the genome sequence accuracy, up until August, 2020, we were unable to access the sequences from Liu et al. *PLoS Pathogens* and Xiao et al. *Nature* that were used to fill the numerous gaps in their genomes. This prevents independent reconstruction of these genomes. Briefly, each group first assembled their genome by pooling sequences across multiple pangolin samples (**Supplementary Table 1**). Xiao et al. justified this approach by explaining that there was only one ‘type’ of coronavirus among the pangolin samples.^2^ Yet, in their study, Xiao et al. isolated particles of the pangolin CoV in Vero E6 culture, and verified the virions by RT-PCR of the Spike and RdRp genes. We propose that it could have been more useful to perform genome or amplicon sequencing on the isolated virus particles, instead of multiple frozen pangolin samples, to fill the gaps in the genome sequence with greater confidence. Similarly, Liu et al. *PLoS Pathogens* pooled sequences from three pangolins from the Guangdong Provincial Wildlife Rescue Center: two of 21 pangolins confiscated in March, 2019 as described in Liu et al. *Viruses*,^4^ and one of six pangolins confiscated in July, 2019.1 Liu et al. *PLoS Pathogens* commented that the July sequences were less abundant than the March sequences, and have not made these sequences available. We could not find a pangolin CoV genome dataset with the accession 2312773 as was provided in the Liu et al. *PLoS Pathogens* Data Availability statement (published on May 14, 2020). We have queried NCBI databases as well as the China National GeneBank Database (https://db.cngb.org/). It is ultimately unclear which parts of the Liu et al. pangolin CoV genome MP789 drew from the July pangolin. Nonetheless, due to their reliance on the same dataset and samples, the Xiao et al. genome GD_1 (GISAID: EPI_ISL_410721) and the Liu et al. genome MP789 (the version that was updated on May 18, 2020) share 99.95% nucleotide identity.

Based on our analysis, we emphasize that there has only been one confirmed source of pangolin CoVs with a Spike RBD nearly identical to that of SARS-CoV-2: pangolins confiscated from smugglers in Guangdong province in March, 2019 (**Fig 2**). The Guangdong pangolin CoV analyses by Zhang et al. and Lam et al. also used the Liu et al. *Viruses* 2019 dataset, and this was clearly stated in their publications.^3,5^ Lam et al. obtained additional March, 2019 smuggled pangolin samples from the Guangzhou Customs Technology Center (NCBI SRA BioProject: PRJNA606875), but only recovered CoV sequences from a single scale sample (**Fig 1A**) and combined this with the Liu et al. *Viruses* dataset to generate their Guangdong pangolin CoV reference.^5^ If there are batches of Guangdong pangolins other than the smuggled pangolins from March, 2019 that have resulted in similar CoV sequences, particularly at the Spike RBD, we have not been able to locate such data based on the Liu et al. and Xiao et al. publications. In comparison to the Guangdong pangolin CoV, the Guangxi pangolin CoVs described by Lam et al. are, once again, less similar to SARS-CoV-2 in the Spike RBD at both the amino acid and nucleotide level even relative to the bat CoV RaTG13 (**Fig 2**). Of the five critical residues for binding between the SARS-CoV Spike RBD and human ACE2 protein, the bat CoV RaTG13 and the Guangxi pangolin CoVs each share only one residue with SARS-CoV-2, whereas the Guangdong pangolin CoV shares five residues with SARS-CoV-2.^5^

**Figure 2.**
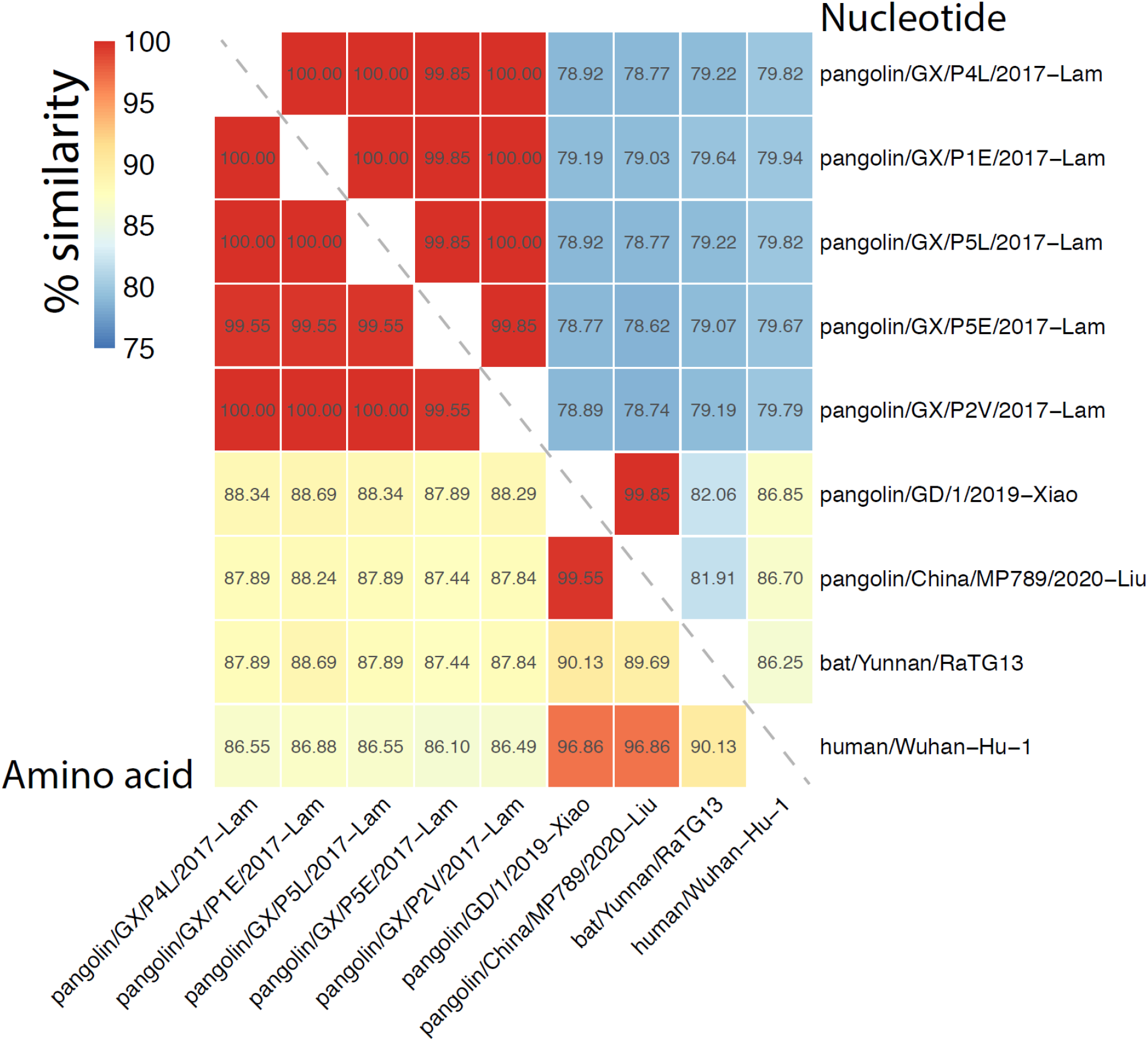
Spike receptor binding domain identity across the pangolin CoVs, bat CoV RaTG13, and human Wuhan-Hu-1 SARS-CoV-2 isolate. The MP789 sequence analyzed here is from the May 18, 2020 update (MT121216.1) although both Xiao et al. and Liu et al. use data published in Liu et al. *Viruses*, 2019. In order to distinguish the updated MP789 from the original MP789, the updated version is indicated as pangolin/China/MP789/<u>2020</u>-Liu instead of pangolin/China/MP789/<u>2019</u>-Liu. Abbreviations: Guangdong, GD; Guangxi, GX.

We would like to reiterate the observations from the literature that it remains to be determined whether the Guangdong and Guangxi pangolin CoVs are found in pangolins in the wild.^1–3,5^ All of the CoV-positive pangolins were confiscated from smugglers, and the Guangdong pangolins were described to have displayed severe respiratory illness and high mortality (14 of the 17 CoV-positive pangolins died within a 1.5 month interval).^2,4,5^ Xiao et al. did not detect CoVs in a second batch of pangolins (four Malayan and four Chinese pangolins) from the Guangdong Provincial Wildlife Rescue Center in August, 2019.^2^ Notably, we do not know how many other batches of smuggled pangolins in China were tested by different groups, found to be CoV-negative, and not published as negative results. In parallel, we do not know what other species of smuggled animals have been tested for SARS-CoV-2. These could suggest a sampling bias that makes it difficult to interpret whether smuggled pangolins are more likely to carry SARS-like CoVs as compared to other species of smuggled animals. Furthermore, a longitudinal study of 334 pangolins confiscated from smugglers in Malaysia, between 2009 to 2019, found that none of the pangolins tested positive for coronavirus. The authors hypothesized that pangolins were an incidental host of coronaviruses potentially due to exposure to infected humans and other smuggled animals.^8^ In the scenario where only two batches of smuggled pangolins possess these SARS-like CoVs, the CoVs could have come from other species held in captivity given the potentially broad host tropism of the related SARS-CoV-2 spike^9–12^ and the fact that each of the two groups of smuggled pangolins carried a single CoV strain. For instance, a Hong Kong smuggling bust in 2014 discovered 40 boxes of pangolin and masked palm civet meat^13^ (palm civets were an intermediate host of SARS-CoV, 2002-2004). In February, 2019, ~30 tons of pangolin, including 61 live pangolins, alongside bat carcasses were raided in Kota Kinabalu.^14^ Which other species were found in the Guangdong and Guangxi anti-smuggling raids, and could these or the human smugglers be hosts of similar CoVs?

One scientist, who read our preprint, inquired whether the above suggestion means that SARS-like CoVs, most closely related to the pangolin CoVs, were circulating among humans before the outbreak of SARS-CoV-2. This is an important point that needs to be elaborated on, particularly for readers who are not specialists in mechanisms of wildlife pathogen spillover into humans. Following the original SARS epidemic in 2002-2004, 13-40% of the tested asymptomatic animal handlers were positive for SARS-reactive antibodies; in comparison, the portion of the general population carrying SARS-reactive antibodies was significantly lower (only up to 0.2% in asymptomatic household contacts).^15,16^ This has been widely interpreted to mean that animal handlers/traders (we include wildlife traffickers in this group) are frequently exposed to wild animals and their pathogens on a regular or prolonged basis. Therefore, it is not surprising that wildlife smugglers would be exposed to novel pathogens and serve as hosts of these pathogens from time to time, without necessarily requiring or resulting in the widespread transmission of these pathogens in the general human population.

In conclusion, our analysis points out that the two complete pangolin CoV genomes, GD_1 by Xiao et al. *Nature*^2^ and MP789 by Liu et al. *PLoS Pathogen*^1^, were built primarily using the same metagenomic dataset published by Liu et al. *Viruses* in 2019. Although there is only a single source of pangolin CoVs that share a near identical Spike RBD with SARS-CoV-2, and there is as yet no direct evidence of pangolins being an intermediate host of SARS-CoV-2, we would like to reinforce that pangolins and other trafficked animals should continue to be considered as carriers of infectious viruses with the potential to transmit into humans.^1–3,5^

**Supplementary Figure 1.**
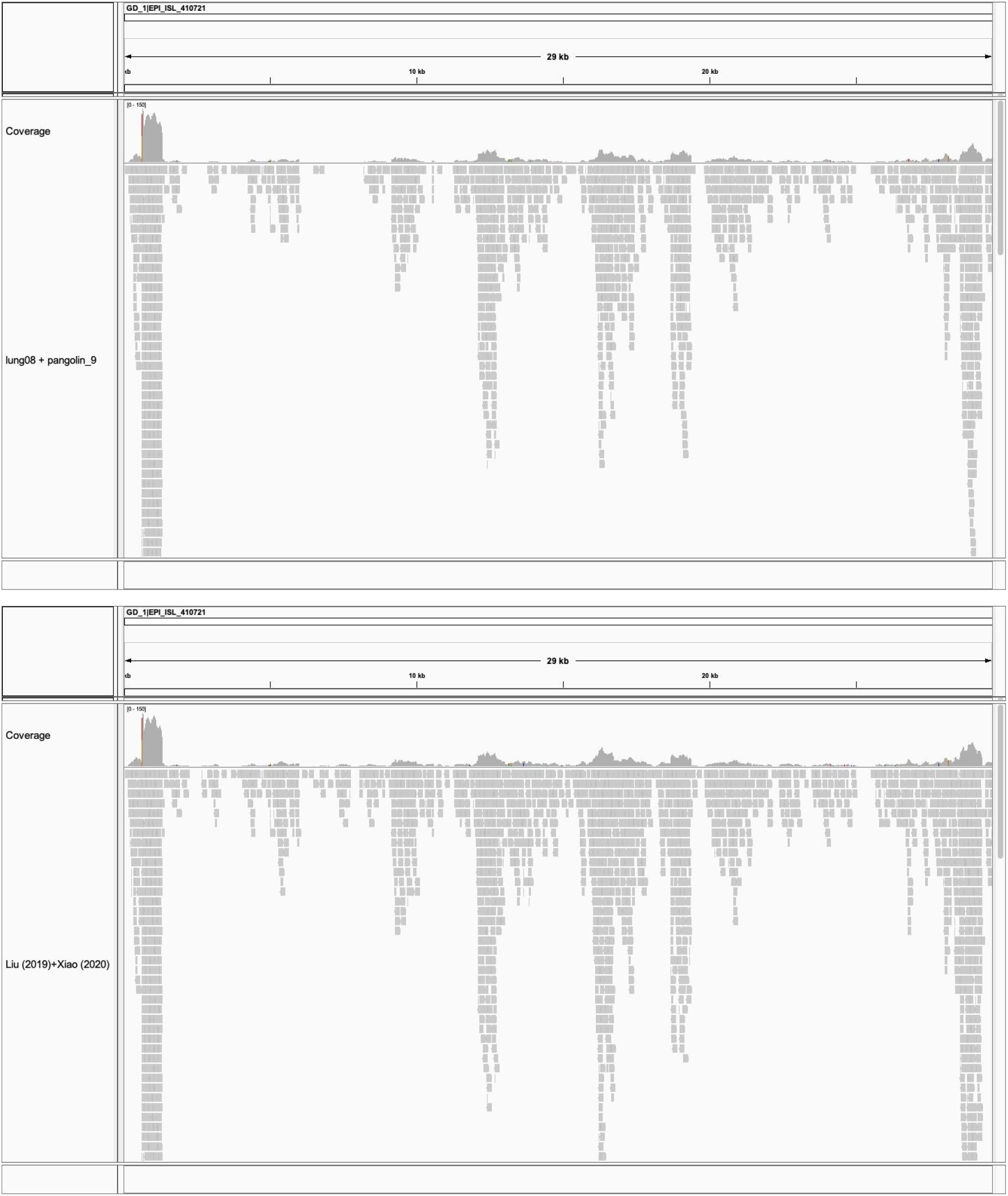
**(Top)** Combined read coverage of the samples lung08 (Liu et al. *Viruses* or M4 in Xiao et al. *Nature*) and pangolin_9 (“M1”, Xiao et al. *Nature*). These two samples contain the highest number of reads mapped to the pangolin CoV genome GD_1 in their respective datasets, and, when pooled, they result in a profile nearly identical to that shown in Xiao et al.’s Extended Figure 4. (Bottom) The two samples dominate the combined profile of all samples from Liu et al. *Viruses* and Xiao et al. *Nature*.

## Materials and Methods

### Read processing and alignment

Raw reads were downloaded from the NCBI SRA (see **Data Availability**). We analyzed 21 and seven paired-end libraries from Liu et al. *Viruses* and Xiao et al. *Nature*, respectively, as well as one single-end library from Lam et al. *Nature* (the scale sample P2S). The reads were filtered and cleaned using *fastp* version 0.20.0 on default settings (trimming adapter sequences, removing poor-quality reads, etc.).^17^ The clean reads were mapped to the genome sequence of GD_1 (GISAID: EPI_ISL_410721) using *minimap2* version 2.17 on default settings.^18^ Duplicate reads were marked and clean reads were coordinate-sorted using *samtools* version 1.10 (sub-commands *markdup* and *sort*, respectively).^19^

### Read coverage statistics

We computed the following read coverage statistics using *bedtools* version 2.29.2^20^: (1) percentage breadth of coverage with respect to GD_1; (2) mean depth of coverage with all mapped reads; and (3) mean depth of coverage without duplicate reads. Also, the per-position read depth data across the GD_1 genome were exported for plotting in R (R Core Team 2020).

### Sequence analysis of the Spike gene

We compared the nucleotide and translated amino acid sequences of the receptor binding domain (RBD) of the Spike gene of the pangolin coronaviruses from Lam et al. *Nature*, Liu et al. *Viruses*, and Xiao et al. *Nature* with the sequences of the bat coronavirus RaTG13 and human coronavirus Wuhan-Hu-1 (their GISAID accessions are provided in **Supplementary Table 2**). We extracted the RBD sequences from GD_1 according to the genome coordinates 22,483-23,151; the receptor binding motif (RBM) sequence embedded within the RBD is located at the genome coordinates 22,837-23,052. The percentage sequence identities were calculated based on pairwise alignments while excluding gapped sites and sites with ambiguous characters. The pairwise sequence similarity matrix was plotted as a heatmap in R^21^.

### Read alignment visualization

The libraries lung08 (from Liu et al. *Viruses*) and pangolin_9 (from Xiao et al. *Nature*) contain the highest number of reads mapped to GD_1 in their respective datasets (**Supplementary Table 1**). We merged the read alignments from those two libraries together for visualization. For the same purpose, we merged all the read alignments from the libraries of Liu et al. *Viruses* and all the read alignments from the libraries of Xiao et al. *Nature*. Read coverage profiles and read alignments were visualized using IGV version 2.8.2.^22^

## Data Availability

The Liu et al. *Viruses* data can be found at NCBI SRA BioProject PRJNA573298 (also accession: SRP223042) and Genome Warehouse BioProject PRJCA002224 (also accession: GWHABKW00000000; https://bigd.big.ac.cn/). The Liu et al. *PLoS Pathogen* pangolin CoV genome sequence (MP789) can be found at NCBI GenBank MT121216 (revised on May 18, 2020, older version: MT084071). The Xiao et al. data can be found at NCBI SRA BioProject PRJNA607174. The Lam et al. data can be found at NCBI SRA BioProject PRJNA606875.

## Acknowledgements

We thank the contributors of the CoV genomes on GISAID (**Supplementary Table 2**). We especially thank Benjamin E. Deverman, Sarah P. Otto, Matthew M. Solomonson, Edwin Cartlidge, and Richard A. White III for reviewing our manuscript. We thank Lamia Wahba for valuable discussions.

## Author contributions

YAC and SHZ contributed equally to this work. YAC: conceptualization, investigation, visualization, writing - original draft. SHZ: conceptualization, data curation, formal analysis, investigation, methodology, software, visualization, writing - review and editing.

## References

1. Liu, P. et al. Are pangolins the intermediate host of the 2019 novel coronavirus (SARS-CoV-2)? PLoS Pathog. 16, e1008421 (2020).

2. Xiao, K. et al. Isolation of SARS-CoV-2-related coronavirus from Malayan pangolins. Nature (2020) doi:10.1038/s41586-020-2313-x.

3. Zhang, T., Wu, Q. & Zhang, Z. Probable Pangolin Origin of SARS-CoV-2 Associated with the COVID-19 Outbreak. Curr. Biol. (2020) doi:10.1016/j.cub.2020.03.022.

4. Liu, P., Chen, W. & Chen, J.-P. Viral Metagenomics Revealed Sendai Virus and Coronavirus Infection of Malayan Pangolins (Manis javanica). Viruses 11, (2019).

5. Lam, T. T.-Y. et al. Identifying SARS-CoV-2 related coronaviruses in Malayan pangolins. Nature (2020) doi:10.1038/s41586-020-2169-0.

6. pangolin 10 - SRA - NCBI. https://www.ncbi.nlm.nih.gov/sra/SRX8582289[accn].

7. Li, X. et al. Pathogenicity, tissue tropism and potential vertical transmission of SARSr-CoV-2 in Malayan pangolins. 2020.06.22.164442 (2020) doi:10.1101/2020.06.22.164442.

8. Lee, J. et al. No evidence of coronaviruses or other potentially zoonotic viruses in Sunda pangolins (Manis javanica) entering the wildlife trade via Malaysia. bioRxiv 2020.06.19.158717 (2020) doi:10.1101/2020.06.19.158717.

9. Damas, J. et al. Broad Host Range of SARS-CoV-2 Predicted by Comparative and Structural Analysis of ACE2 in Vertebrates. bioRxiv 2020.04.16.045302 (2020) doi:10.1101/2020.04.16.045302.

10. Qiu, Y. et al. Predicting the angiotensin converting enzyme 2 (ACE2) utilizing capability as the receptor of SARS-CoV-2. Microbes Infect. (2020) doi:10.1016/j.micinf.2020.03.003.

11. Frank, H. K., Enard, D. & Boyd, S. D. Exceptional diversity and selection pressure on SARS-CoV and SARS-CoV-2 host receptor in bats compared to other mammals. bioRxiv 2020.04.20.051656 (2020) doi:10.1101/2020.04.20.051656.

12. Johnson, B. A. et al. Furin Cleavage Site Is Key to SARS-CoV-2 Pathogenesis. 2020.08.26.268854 (2020) doi:10.1101/2020.08.26.268854.

13. Hong Kong Customs and Excise Department - Press Releases. https://www.customs.gov.hk/en/publication_press/press/index_id_1221.html (2010).

14. Record setting 30-tonne pangolin seizure in Sabah ahead of World Pangolin Day - Wildlife Trade News from TRAFFIC. https://www.traffic.org/news/record-setting-30-tonne-pangolin-seizure-in-sabah-ahead-of-world-pangolin-day/.

15. Cyranoski, D. & Abbott, A. Virus detectives seek source of SARS in China&#x2019;s wild animals. Nature 423, 467 (2003).

16. Cheng, V. C. C., Lau, S. K. P., Woo, P. C. Y. & Yuen, K. Y. Severe acute respiratory syndrome coronavirus as an agent of emerging and reemerging infection. Clin. Microbiol. Rev. 20, 660–694 (2007).

17. Chen, S., Zhou, Y., Chen, Y. & Gu, J. fastp: an ultra-fast all-in-one FASTQ preprocessor. Bioinformatics 34, i884–i890 (2018).

18. Li, H. Minimap2: pairwise alignment for nucleotide sequences. Bioinformatics 34, 3094–3100 (2018).

19. Li, H. et al. The Sequence Alignment/Map format and SAMtools. Bioinformatics 25, 2078–2079 (2009).

20. Quinlan, A. R. & Hall, I. M. BEDTools: a flexible suite of utilities for comparing genomic features. Bioinformatics 26, 841–842 (2010).

21. Ripley, B. D. The R project in statistical computing. MSOR Connections. The newsletter of the LTSN Maths (2001).

22. Thorvaldsdóttir, H., Robinson, J. T. & Mesirov, J. P. Integrative Genomics Viewer (IGV): high-performance genomics data visualization and exploration. Brief. Bioinform. 14, 178–192 (2013).

